# Stress in utero: effects on gliovascular integrity in female infant offspring

**DOI:** 10.1101/2024.04.17.589633

**Authors:** Magda Rodrigues, Inês Sousa, Filipa Baptista, Vanessa Coelho-Santos

## Abstract

From early in life, experiences like prenatal stress profoundly affect long-term health and behavior. Maternal stress increases fetal exposure to glucocorticoids (GC), disrupting neurodevelopment and raising susceptibility to psychiatric disorders. Previous studies on synthetic GCs, such as dexamethasone (DEX), revealed impairments in neurogenesis and dendritic spine development. However, the influence of prenatal stress on the gliovascular interface remains unclear. This interface, involving the relationship between astrocytes and blood vessels, is essential for healthy brain development. Our study demonstrates that prenatal stress alters the expression and localization of astrocytic proteins crucial for maintaining vascular homeostasis, such as aquaporin-4, in female offspring exposed to DEX. While overall vascular density remains unaffected, it triggers morphological changes. Particularly, the hippocampus and prefrontal cortex exhibit heightened vulnerability to these effects. This study reveals prenatal stress as a potent disruptor of gliovascular development, urging deeper inquiry into its implications.

## Introduction

Early-life experiences, like prenatal stress, can deeply shape long-term health and behavioural patterns (reviewed in (Bock et al., 2015)). Prenatal stress is often linked to increased levels of glucocorticoids (GCs), which can arise from maternal stress or the administration of GC therapy during gestation. This therapy is utilized to manage women at risk of preterm labor or to treat fetuses at risk of congenital adrenal hyperplasia. Such prenatal stress significantly impacts offspring mental health, as discussed in the review article (Krontira et al., 2020). Maternal GCs can be transmitted to the developing fetus through the placenta (Krontira et al., 2020), where elevated levels during critical periods of development disrupt neurodevelopmental processes, increasing offspring susceptibility to psychiatric disorders by triggering structural and functional abnormalities in the developing brain (Davis et al., 2013; Drozdowicz & Bostwick, 2014; Khulan & Drake, 2012). The mental health challenges in adult offspring include schizophrenia-related behaviors in males and anxious-depressive like symptoms in females, alongside with social deficits increased emotional reactivity and drug-seeking behaviors (Borges et al., 2013; Caetano et al., 2017; Oliveira et al., 2006, 2012; Rim et al., 2022; Rodrigues et al., 2012).

Several structural brain alterations have been reported upon prenatal DEX exposure, namely a decrease in the number of neuronal cells and volumetric atrophy of the nucleus accumbens (Rodrigues et al., 2012), and impaired radial migration of post-mitotic neurons during the development of the cerebral cortex (Fukumoto et al., 2009). Prenatal exposure to DEX increases the volume of the bed nucleus of the stria terminalis, whereas it reduces the volume of the amygdala due to dendritic atrophy (Oliveira et al., 2012). At neuronal levels, prenatal exposure GC namely to dexamethasone (DEX, a synthetic GC) has been found to impair neurogenesis (Tsiarli et al., 2017) and dendritic spine development (Oliveira et al., 2012; Tanokashira et al., 2012), compromise neuronal survival (Crochemore et al., 2005) and lead to morphological alterations in axons and dendrites of developing neurons (Pinheiro et al., 2018). Prenatal DEX also affects microglia, the immune cells of the central nervous system, that seems to be indicated to explain the sex-biased onset and vulnerability to psychiatric disorders (Caetano et al., 2017; Gaspar et al., 2021; Rim et al., 2022).

The brain vasculature appears to be also susceptible to the effects of stress (Dion-Albert et al., 2022; (reviewed in (Welcome & Mastorakis, 2020) including compromises the integrity of the blood-brain barrier (BBB), which plays a crucial role in regulating the passage of molecules and cells between the bloodstream and the brain. Both vasculature and its barrier integrity are vital for brain development and function (Coelho-Santos & Shih, 2020). Disruptions in the BBB during critical developmental periods can have long-lasting consequences on neuronal connectivity, synaptic plasticity, and overall brain function (Ouellette et al., 2020), and a predisposition to psychiatric disorders (Reviewed in (Dion-Albert et al., 2023)). Alterations in the vasculature and the BBB integrity have been demonstrated to play a role in adult stress responses, influencing both resilience and vulnerability to psychiatric disorders (Dion-Albert et al., 2022; Dudek et al., 2020; Najjar et al., 2017). Despite the scarcity of research on prenatal stress, emerging evidence hints at its potential influence on brain vasculature. Rats exposed to prenatal DEX (subcutaneous injection 100 mg/kg on G14-21) present reduced vascularization (vascular area fraction) in the CA3 region of the hippocampus at adulthood (P90) comparing to controls (Neigh et al., 2010). In a mouse model of prenatal DEX exposure, it was found that at P4 there is a significant downregulation of the tight junction protein claudin-5 in total brain samples after multiple intraperitoneal injections with DEX (0.1 mg/kg at G15-17). Immunofluorescence images of claudin-5 and of the endothelial marker Pecam-1/CD31 revealed changes in vessel morphology (Neuhaus et al., 2015). Previous study in sheep showed that single maternal treatment with DEX (four 6 mg intramuscular injections of dexamethasone every 12 h over 48 h) lead to an increase of claudin-5 in the cerebral cortex and upregulation of occludin in the cerebellum. Multiple exposures (once a week for 5 consecutive weeks) did not impact on claudin-5 expression levels in the cerebellum but upregulated occludin in the cerebral cortex and cerebellum (Sadowska et al., 2010), suggesting that changes in the brain are region-specific.

Interestingly, the effects of prenatal stress on the gliovascular interface remain elusive and require further clarification. The gliovascular interface, where astrocytes intimately interact with blood vessels in the brain, is a critical hub for maintaining the integrity of the BBB (Gilbert et al., 2019). This dynamic interface has been also described as participant in the neurovascular coupling (Freitas-Andrade et al., 2023), ensuring optimal blood flow in response to neuronal activity. Therefore, unraveling the complex interplay between astrocytes and vessels is crucial for understanding the minutiae of healthy brain development. To address this gap and investigate if gliovascular unit alterations are linked to prenatal exposure to stress, we subcutaneously administered physiological saline or 50 µg/kg of DEX to pregnant mice from gestational days 16 to 18, simulating the induction of stress hormones during the prenatal stage. We collected the brains of offspring at postnatal day 14 (P14) and utilized immunohistochemistry techniques to assess gliovascular unit maturation in several brain regions, including the hippocampus, prefrontal and somatosensory cortex, striatum, and cerebellum. This study invites further investigation into how prenatal stress shapes infant gliovascular development, potentially influencing adult behavior.

## Methods

### Ethics statement

Procedures involving animals were approved by the Animal Welfare Committee of the Coimbra Institute for Clinical and Biomedical Research (iCBR), Faculty of Medicine, University of Coimbra (ORBEA 03/2021, and the procedures were performed by licensed users of the Federation of Laboratory Animal Science Associations (FELASA), and accordingly with the guidelines of the European Parliament (2010/63/EU), translated to the Portuguese law in 2013 (Decreto-lei nº 113/2013).

### Animals and Experimental Design

Female C57BL/6J mice weighing 19-25 g at reproductive age were housed in certified facilities, with a temperature and humidity-controlled environment, under a 12h:12h light-dark cycle and with *ad libitum* access to water and food.

Female mice were mated with males (1 male per female) for a period of 24 h. Male bedding was added to female’s cages the day prior male presentation. Gestational day 0 (GD0) was considered the day after mating day.

Aiming to achieve behavioral alterations induced by prenatal stress (namely depressive-like behavior in females), the experimental model described by (Rim et al., 2022) was adopted. Dams were arbitrarily assigned to control (CTR) or dexamethasone (DEX) experimental groups. DEX at the dose of 50 μg/kg or saline (vehicle) were administered subcutaneously to pregnant mice on GD16, 17 and 18. This dose mimics mice physiological GC levels induced by a stress condition. Female offspring of CTR or DEX dams were sacrificed at postnatal day 14 (P14). Figure 1 A illustrates a graphical overview of the experimental arrangement.

**Figure 1.**
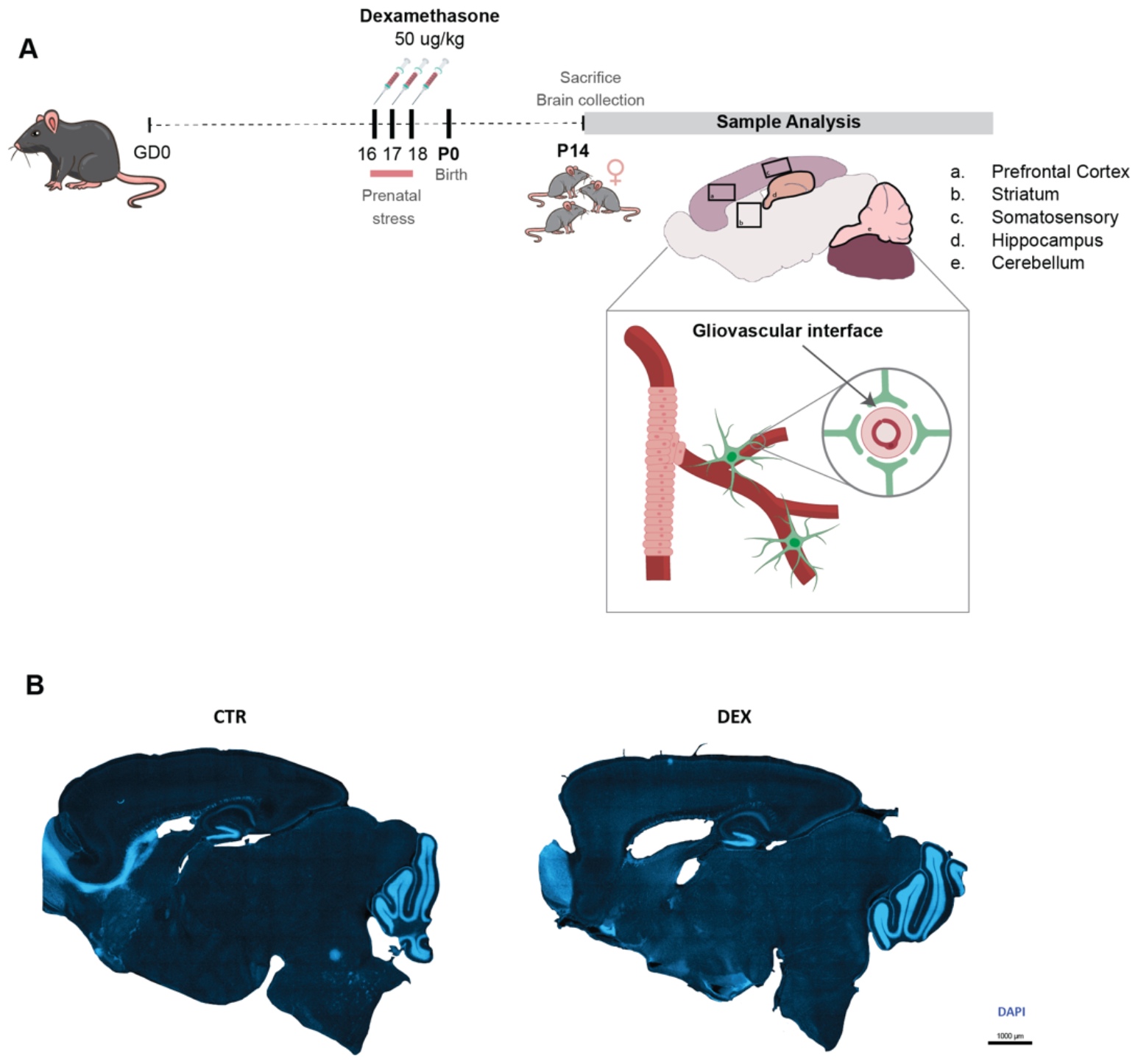
Schematic representation of study methodology. A) This figure provides a visual overview of the research plan executed, outlining the key methods employed and the overarching objectives of the study. B) This panel presents microscopic images showcasing brain tissue sections from the two experimental groups: control (CTR) and dexamethasone (Dex)-treated. Images captured using slide scanner, identification of cell nuclei using DAPI (blue).

### Sample collection

At P14, female offspring mice were deeply anesthetized with an intraperitoneal injection of ketamine (50mg/kg; Nimatek, Dechra, UK) and xylazine (2mg/kg; Sedaxylan, Dechra, UK). Animals were then transcardially perfused with 0.1M phosphate-buffered saline (PBS, in mM: 137 NaCl, 2.7 KCl, 10 Na2HPO4, 1.8 KH2PO4, pH 7.4), followed by 4% paraformaldehyde (PFA) in 1% PBS. Brains were postfixed in 4% PFA, transferred to a solution of 30% sucrose in PBS-1% and stored at 80°C until processing.

### Free-floating immunohistochemistry

Sagittal sections (Figure 1B, 100 μm) were obtained using a cryostat (Leica CM3050S) and submerged in PBS containing 0.01% sodium azide (Thermo fisher, 26628-22-8), then stored at 4ºC until further use. Prior to immunostaining, slices underwent a 2h incubation at room temperature (RT) in a blocking solution consisting of 0.1M PBS supplemented with 5% (v/v) goat serum (Sigma; G9023) and 0.3% (v/v) Triton X-100 (Fisher Scientific, BP151-100). Subsequently, slices were transferred to well plates containing primary antibodies (refer to Table 1) diluted in 0.1M PBS with 5% (v/v) goat serum and 0.2% (v/v) Triton X-100 and incubated at 4ºC for 48 hours. Following this, slices underwent triple rinsing for 15 min each with 0.1M PBS, succeeded by three 5 min rinses with 0.1M PBS, under gentle agitation. For specimens treated with nonconjugated primary antibodies, secondary antibodies (refer to Table 1), prepared in PBS supplemented with 5% (v/v) goat serum and 0.2% (v/v) Triton X-100, were applied and incubated for 2h at RT, followed by repeated washes. Post-immunostaining, slices were mounted, air-dried, and sealed with DAPI Fluoromount-G (Invitrogen, 00-4959-52) prior to imaging. Whole slide imaging was acquired using a Slide scanner (Carl Zeiss Axio Scan.Z1) equipped with a 20x objective (Plan-Apochromat 20x/0.8 M27), capturing the maximum z-stack images feasible. High-resolution confocal images were acquired using a Confocal microscope (Zeiss Confocal Microscope LSM 710, Axio Examiner) with a 40x objective lens (oil-immersion, Plan-Apochromat 40x/1.4 Oil DIC M27), capturing images with dimensions of 1024 x 1024 pixels and averaging performed for every 2 frames.

**Table 1.**

|  | Dilution | Fluorophore | Host | Supplier (Cat No) | Target |
| --- | --- | --- | --- | --- | --- |
| Primary Antibodies |  |  |  |  |  |
| Lectin | 1:250 | Fluorescein | *Lycopersicon esculentum* | Vector Labs, FL-1171-1 | Endothelial cells |
| Aquaporin 4 | 1:250 | CoraLite 647 | Rabbit | Proteintech, CL647-16473 | Astrocytes endfeet |
| Secondary antibodies |  |  |  |  |  |
| Donkey anti-Mouse | 1:250 | Alexa Fluor 555 | Donkey | Invitrogen, A-31570 |  |

### Analysis of histological data

#### Aquaporin 4 expression analysis

In our study, we investigated the colocalization of aquaporin 4 (AQ4) with lectin (Lec) to demonstrate the association between astrocyte endfeet and blood vessels. Utilizing the ImageJ (Fiji), fluorescence images obtained from the Slide Scanner were converted into 2D images through max intensity z-projection. Subsequently, a region of interest (ROI) was delineated, covering most of the brain regions, from prefrontal, striatum, somatosensory cortex, hippocampus and cerebellum. Colocalization analysis of fluorescence microscopy images was conducted using BIOP’s (Bioimaging and Optics Platform) version of JACoP (Just Another Colocalization Plugin) (Bolte & Cordelières, 2006). The assessment involved measuring Pearson’s Correlation Coefficient. Pearson’s correlation coefficient was highlighted as it quantifies the strength and direction of the linear relationship between two variables, specifically the image pixels of fluorescence, providing insights into the colocalization patterns of AQ4 and Lec in astrocyte endfeet and vessels.

The total expression of AQ4 was quantified within the predefined ROIs. Mean gray value and area measurements of the AQ4 channel were obtained from these ROIs. Background subtraction was performed by averaging the mean gray values from two background regions. Then, background value was subtracted from the mean gray value of AQ4. Image volume was calculated by multiplying the ROI area by the number of z-stacks used in each image (mm^3^). Finally, to calculate the total AQ4 expression the mean gray value obtained was divided by the volume (Fluorescence AU/mm3).

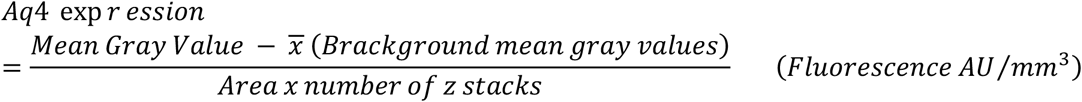

#### Vasculature analysis

To assess the impact of DEX on the vasculature, small ROIs were isolated from the previously described areas, utilizing the Lec channel. Prior to analysis, images were prepared by adjusting brightness and contrast to enhance visualization, followed by conversion into grayscale. Vessel tracing was meticulously conducted using the hand-free tracing tool in ImageJ, facilitating the creation of overlays on the z-stack images of vessels. Subsequently, various parameters were quantified to evaluate vascular morphology. The length of vessels was measured, while the number of branches was determined by recording the newly added traces during vessel tracing. Branching points were carefully annotated whenever observed during the tracing process. In addition to quantitative assessments, a qualitative analysis of vessel tortuosity was performed. Images were categorized based on the degree of tortuosity observed, with three distinct levels identified: “None,” indicating no visible alterations in vessel morphology; “Low,” denoting the presence of less than five vessels exhibiting tortuosity; and “High,” indicating five or more vessels displaying noticeable roughness or tortuosity. Finally, the results were presented based on the volume of each image (Area x number of z stacks).

#### Exclusion criteria

From the slide scanner, z-stacks that exhibited blurriness in the images were excluded from further analysis.

### Statistics

Data analysis was conducted using GraphPad Prism 10 software. Graphical representations show mean values accompanied by the standard error of the mean (SEM). Tests of normality were first performed to validate the use of parametric tests. The Shapiro-Wilk normality test was employed to assess the normal distribution of the data. Parametric tests were applied to normally distributed data, while non-parametric tests were utilized for non-Gaussian distributed data. Statistical significance was defined as a P value of < 0.05. Further details regarding statistical tests are provided in the figure legends.

## Results

### Maternal stress effects on development of the cerebrovascular network

Prenatal stress, a critical environmental factor, can leave lasting scars on the central nervous system (CNS), potentially leading to long-term functional impairments in offspring and increasing susceptibility to neuropsychiatric disorders later in life (Borges et al., 2013; Caetano et al., 2017; Oliveira et al., 2006, 2012; Rim et al., 2022; Rodrigues et al., 2012). Recognizing its impact on infancy is essential for early intervention strategies. With this in mind and to gain insight into the blood vessels network within the brain of infant female mice, we used immunohistochemistry to stain vessels with lectin. This method offers a clear visualization of the brain vasculature, making it suitable for quantitative analysis. We were able to analyze and characterize the 3D morphology of the vascular network in detail. The key advantage of our free-hand method lies in its ability to facilitate quantitative analysis of the vasculature (Figure 2A). This means we can not only visualize the vessels but also measure specific features to gain deeper insights into their structure.

**Figure 2.**
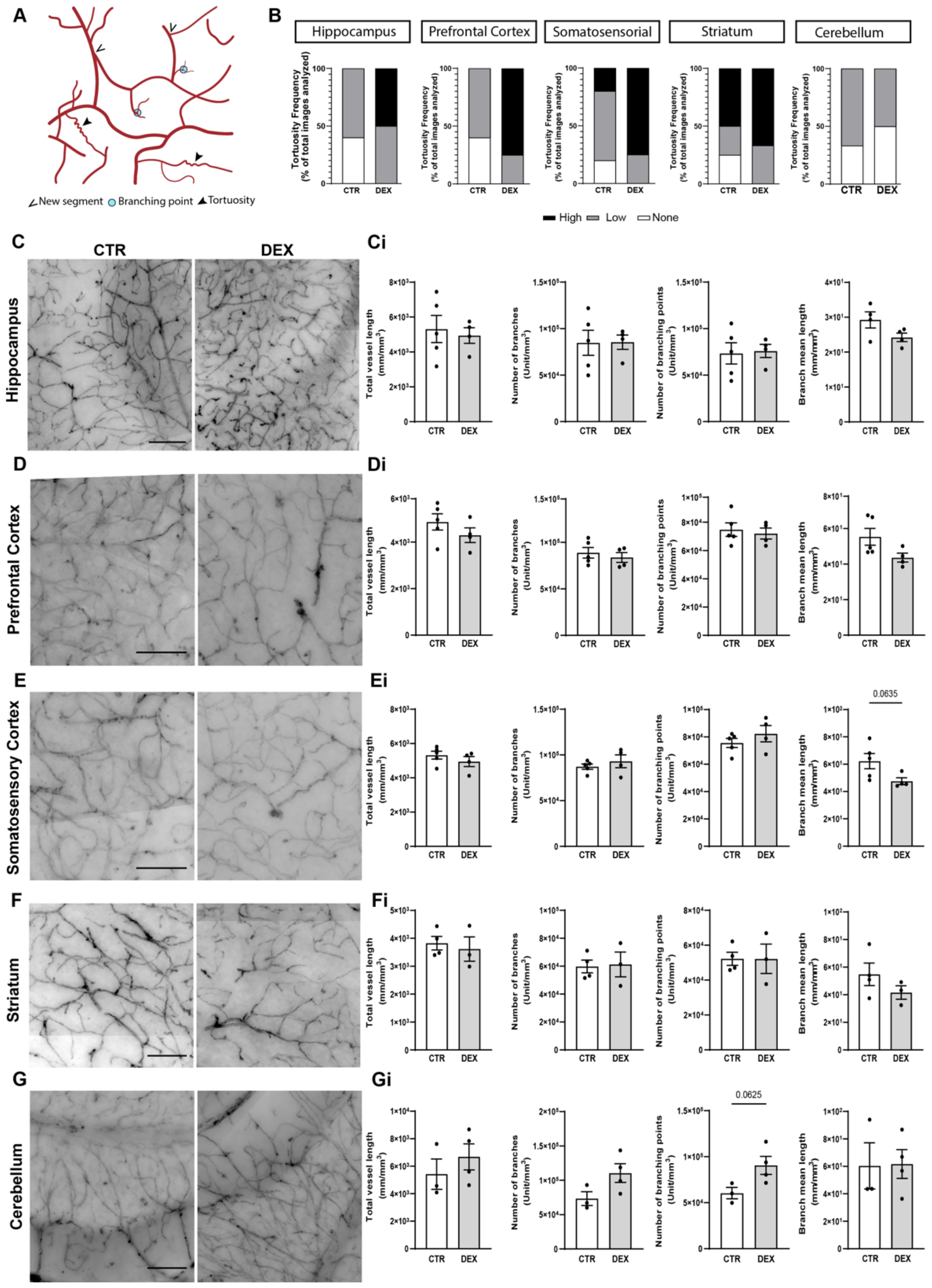
Impact on brain vasculature in postnatal day 14 female offspring exposed to maternal stress. A) Schematic representation of the analysis conducted. B) Qualitative analysis of vessel tortuosity observed in microvessels, revealing an overall increase in all brain regions studied of animals exposed to maternal stress. C-G) Overview of brain vasculature, representative pictures of vasculature in the selected brain regions, hippocampus (C), prefrontal and somatosensory cortex (D,E), striatum (F) and cerebellum (G). Results from this analysis are displayed as total vessels length, number of branches, branching points and branches mean length (Ci-Gi). No significant alterations were observed in vessel density or vessel branching in the brain regions of female offspring. Scale bar 100 µm. Results are presented as the mean ± SEM, with n=3-5 animals. Results compared to the control, as determined by the Mann-Whitney test.

Both CTR and DEX-exposed groups exhibited some degree of vessel tortuosity. Intriguingly, our analysis revealed a notable increase in highly tortuous vessels in most of the analyzed brain regions in female’s offspring exposed to prenatal stress (Figure 2B). Compared to CTR, the hippocampus displayed a 50% increase in vessels with high tortuosity. Similarly, the prefrontal and somatosensory cortex exhibited increases of 75% and 55%, respectively, in highly tortuous vessels. While the striatum also showed an increase, it was more modest, with only a 16.17% rise compared to CTR. Interestingly, the cerebellum was the only region where we did not observe noteworthy changes in tortuosity.

Overall, the vascular density (number of vessels per unit volume) remained largely unaffected by prenatal stress. Because branching points are crucial for creating a complex vascular network, allowing for efficient blood flow distribution, we delved deeply into your analysis. However, although not statistically significant, a trend towards a decrease in the overall length of the branches was observed in the hippocampus (CTR 29.24±2.313 mm/mm^3^, DEX 24.23±1.323 mm/mm^3^), prefrontal cortex (CTR 55.36±4.763 mm/mm^3^, DEX 43.65±2.497 mm/mm^3^), somatosensory cortex (CTR 62.21±5.605 mm/mm^3^, DEX 47.56±2.505 mm/mm^3^, p=0.0635), striatum (CTR 54.77±8.146 mm/mm^3^, DEX 41.62±4.887 mm/mm^3^) following prenatal stress exposure (Figure 2Ci-Fi). Conversely, no such trend was observed in the cerebellum (CTR 60.38±16.83 mm/mm^3^, DEX 61.72±10.49 mm/mm^3^, Figure 2Gi). In addition, the number of branches and branching points within the hippocampus (branches CTR 8.474x10^4^±1.339x10^4^ Unit/mm^3^, DEX 8.547x10^4^±7.775x10^3^ Unit/mm^3^, branching points 7.336x10^4^±1.148x10^4^ Unit/mm^3^, DEX 7.596x10^4^±7.071x10^3^ Unit/mm^3^), prefrontal cortex (branches CTR 8.879x10^4^±5.688x10^3^ Unit/mm^3^, DEX 8.975x10^4^±5.319x10^3^ Unit/mm^3^, branching points 7.481x10^4^±4.804x10^3^ Unit/mm^3^, DEX 7.204x10^4^±3.955x10^3^ Unit/mm^3^), somatosensory cortex (branches CTR 8.717x10^4^±2.827x10^3^ Unit/mm^3^, DEX 9.309x10^4^±6.982x10^3^ Unit/mm^3^, branching points 7.554x10^4^±3.311x10^3^ Unit/mm^3^, DEX 8.233x10^4^±5.925x10^3^ Unit/mm^3^) and striatum (branches CTR 5.980x10^4^±4.603x10^3^ Unit/mm^3^, DEX 6.126x10^4^±8.866x10^3^ Unit/mm^3^, branching points 5.220x10^4^±3.759x10^3^ Unit/mm^3^, DEX 5.220x10^4^±8.437x10^3^ Unit/mm^3^) remained largely unchanged across CTR and DEX conditions (Figure 2Ci-Fi). The cerebellum exhibited a trend towards an increase in branching points (CTR 6.0347x10^4^±6.134x10^3^ Unit/mm^3^, DEX 9.0542x10^4^±9.743x10^3^ Unit/mm^3^, p=0.0625, Figure 2Gi).

In sum, our results show that prenatal stress in female offspring increased vessel tortuosity in the hippocampus, prefrontal cortex, somatosensory cortex, and striatum. Additionally, a trend towards shorter segment length was observed in these regions. These findings suggest that prenatal stress may disrupt blood vessel development without causing major vascular architecture, as evidenced by the lack of significant changes in branching patterns.

### Prenatal stress exposure alters the regulation of AQ4 in the brains of female offspring

AQ4, a crucial protein residing in the endfeet of astrocytes, plays a vital role. These extensions tightly wrap around brain microvessels, acting as gatekeepers for water transport and maintaining the delicate homeostasis of fluids within the brain (Salman et al., 2022). While this interface is critical for the development of brain blood vessel maturation (Gilbert et al., 2019), the impact of prenatal stress has not been previously described. Hence, our research focused on investigating the impact of prenatal stress on this complex system. Once again, the prefrontal cortex and hippocampus were the regions most impacted by DEX exposure. In the hippocampus, we observed a significant increase in AQ4 protein expression levels (CTR 1.85x10^5^±1.104x10^4^ AU/mm^3^, DEX 6.013x10^5^±1.416x10^5^ AU/mm^3^, p=0.0264, Figure 3A-Ai) that remained concentered at the vascular area (CTR 0.563±0.030, DEX 0.465±0.098, p=0.379, Figure 3A-Ai). In the prefrontal cortex, we noticed a significant increase in the coverage of blood vessels by AQ4 (CTR 0.452±0.027, DEX 0.617±0.015, p=0.0018, Figure 3B-Bi); however, the total amount of AQ4 protein remained unchanged (CTR 1.192x10^5^±1.717x10^4^ AU/mm^3^, DEX 1.694x10^5^±3.861x10^4^ AU/mm^3^, p=0.279, Figure 3 B-Bi). This suggests a change in the distribution of existing AQ4 in the prefrontal cortex. Interestingly, we showed that the expression and vascular coverage by AQ4 prevailed unaltered at somatosensory cortex (1.893x10^5^±6.541x10^4^ AU/mm^3^, p=0.806, 0.513±0.052, p=0.200, Figure 3C-Ci), striatum (2.231x10^5^±8.417x10^4^ AU/mm^3^, p=0.606, 0.547±0.088, p=0.298, Figure 3D-Di) and cerebellum (6.708x10^5^±3.958x10^5^ AU/mm^3^, p=0.564, 0.420±0.082, p=0.751, Figure 3E-Ei) at the offspring exposed to prenatal stress (DEX) compared to controls.

**Figure 3.**
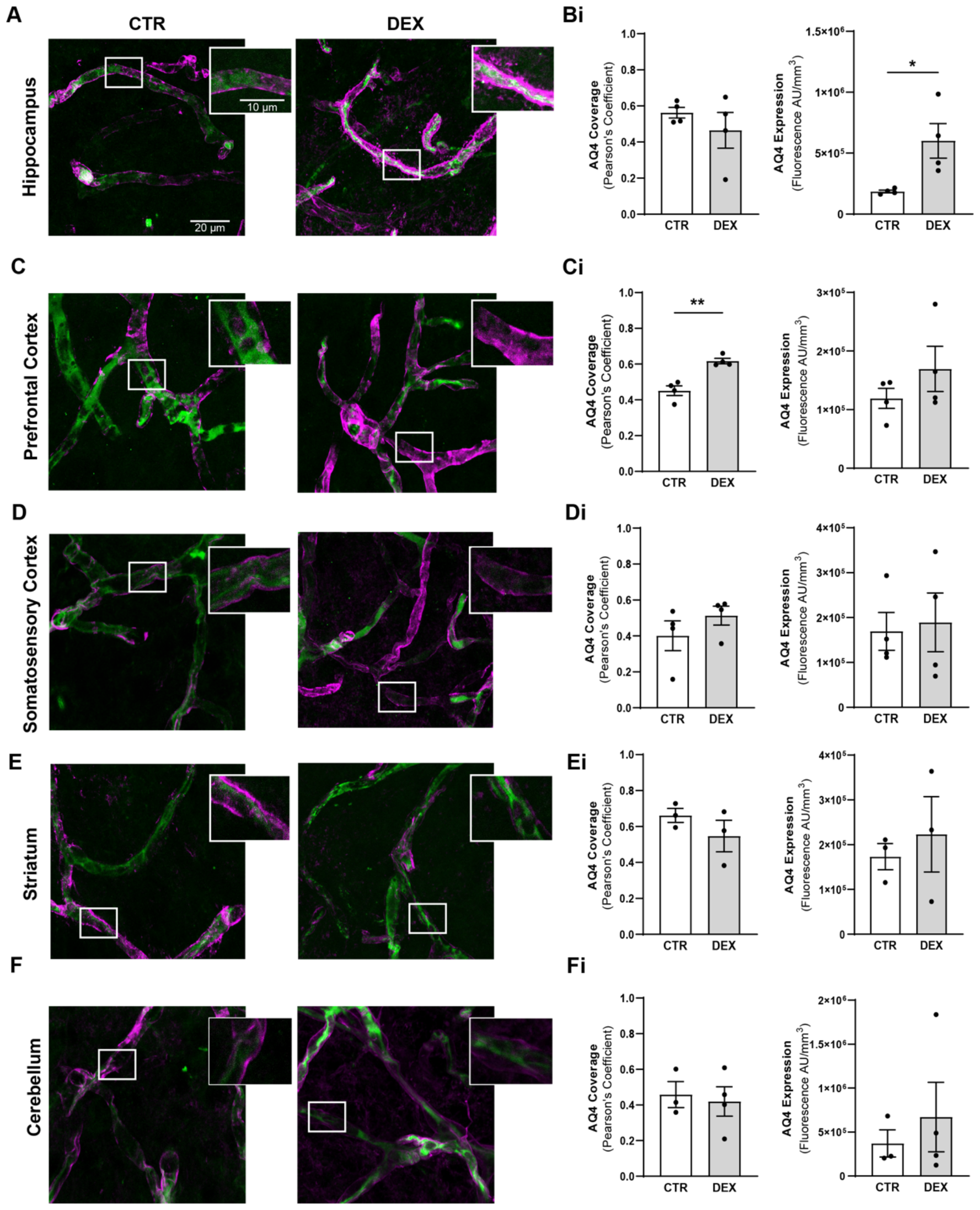
Differential response of aquaporin-4 astrocytic endfeet expression and distribution across brain regions in female mice exposed to maternal DEX. A-G) High-resolution images 60 μm projections show astrocytic endfeet coverage surrounding immunostained microvessels (Lec - green), aquaporin-4 (AQ4) protein (magenta) localization within these endfeet processes across five distinct brain regions of female mice: hippocampus (A), prefrontal cortex (B), somatosensory cortex (C), striatum (D), and cerebellum (E). Ai-Ei) Quantitative data obtained, total AQ4 expression levels and the colocalization between AQ4 and microvessels (calculated using the Pearson coefficient) were analyzed within each brain region. Results are presented as the mean ± SEM, with n=3-5 animals. *p<0.05, **p<0.05 compared to the control, as determined by the unpaired t-test.

Our findings present that prenatal stress disrupts the normal developmental distribution and expression levels of AQ4 in particular at prefrontal cortex and hippocampus and suggest that it might affect their function.

## Discussion

Prenatal stress, a significant early life adversity, has the potential to imprint lasting effects on the CNS, increasing the risk of neuropsychiatric disorders later in life. The release of GC triggered by prenatal stress, potentially impacting the developing vasculature (Millage et al., 2017) and astrocytes proliferation (Vivi & Di Benedetto, 2024) that happens in this frame time. However, its impact on the gliovascular interface remains unexplored. To fill this gap in our understanding, we investigated the effects of elevated glucocorticoids on gliovascular interface development. For this purpose, we employed a prenatal stress model involving exposure to a synthetic GC (DEX). To the best of our knowledge, our findings reveal, for the first time, that prenatal stress, echoing human neuropsychiatric symptoms, notably anxiety and depressive-like behaviors in females later in life, might be associated with changes in the gliovascular interface during infancy.

The limited research conducted thus far indicates that prenatal exposure to DEX may influence brain vasculature, hinting at region-specific alterations within the brain. Because of that we expanded our analysis to include the gliovascular unit in various brain regions: prefrontal cortex, striatum, somatosensory cortex, hippocampus, and cerebellum. Our prenatal stress model increased vessel tortuosity, characterized by a corkscrew-like appearance, in the hippocampus, prefrontal cortex, somatosensory cortex and striatum of female offspring. While there was a trend towards shorter vessel length, overall vascular density and branching patterns remained largely unaffected. This increase in tortuosity may be linked to alterations in the structure of endothelial cells lining the blood vessel walls, which have been reported in other studies investigating glucocorticoid-induced endothelial cell dysfunction (Han, 2012), due to changes in tight junction proteins, which are crucial for maintaining the integrity of the BBB (Neuhaus et al., 2015). Likewise, existing research explores how DEX affects endothelial cell activity, demonstrating increases in their proliferation and migration (Polytarchou & Papadimitriou, 2005; Wallerath et al., 2004). This damage could manifest as weakened vessel structure, further impacting vessel function, which could disrupt blood flow and the homeostasis within these brain regions. In addition, we spotted a trend towards decreased mean branches length in the hippocampus following prenatal DEX exposure. This finding aligns with prior reports (Neigh et al., 2010) on DEX-induced decrease in hippocampal vasculature. We also report a similar tendency for decrease branch segment length in the cortex and striatum following prenatal DEX exposure. Interestingly, these DEX-induced effects appear to be region-specific, as neither a decrease in vascular segment length nor an increase in tortuosity was observed in the cerebellum.

Previous studies have shown that early life stress impacts astrocytic structure and function, particularly astrogliosis markers such as GFAP, GLT, and GLAST, in ways that may have implications later in life (Majcher-Maślanka et al., 2019; Martisova et al., 2013; Orso et al., 2023; Zeng et al., 2020). Although it has been shown that astrocyte plasticity in adults responds to prenatal development (Shende et al., 2015), there is a significant knowledge gap concerning the impact during infancy and whether this constitutes an early effect. Our study revealed no significant changes in the localization or expression levels of the protein AQ4 in the somatosensory cortex, striatum, and cerebellum of female offspring exposed to prenatal stress. However, in the prefrontal cortex, we observed an interesting phenomenon. While overall AQ4 production remained unchanged, there was an increase in AQ4 co-localization with blood vessels, suggesting a potential shift in its location within the tissue. Interestingly, this alteration did not coincide with a reduction in the overall number or length of vessels in the area. In contrast, the hippocampus displayed a different response. Here, prenatal stress led to a significant increase in AQ4 production itself. These findings suggest that prenatal stress may have region-specific effects on astrocytic endfeet regarding its AQ4 function, potentially impacting its function in the prefrontal cortex and hippocampus. AQ4 within astrocytic endfeet plays a critical role in managing water dynamics within the brain. Located at the perivascular space, AQ4 acts as a connector between astrocytes and the extracellular space surrounding blood vessels (Nakada et al., 2017). This allows for the controlled efflux of water from astrocytes, a crucial process for maintaining proper water balance. AQ4 facilitates water movement from the interstitial space back into astrocytes. By controlling water efflux and potentially influx, AQ4 helps maintain fluid balance within the BBB and contributes to the essential flow of cerebrospinal fluid (Haj-Yasein et al., 2012; Igarashi et al., 2014; Nakada et al., 2017). Thus, an excessive increase in AQ4 expression could also lead to disruptions in water balance within the brain. This could potentially contribute to conditions like cerebral edema (fluid buildup in the brain). AQ4 function in adult brains has been well explored, however its role during development remains unclear (Fallier-Becker et al., 2014; Wen et al., 1999). Some believe AQ4 helps reduce brain water content after birth, as brains lacking AQ4 (AQP4-KO mouse) take longer to lose excess water (Li et al., 2013). Additionally, AQ4 appears alongside astrocyte development around blood vessels, suggesting a link to neurovascular unit formation. Studies show AQ4 expression strengthens around blood vessels by P14, coinciding with BBB formation in mice (Lunde et al., 2015). Indeed, AQ4 has been also associated with BBB maturation (Li et al., 2013). Our results suggest that since prenatal stress disrupts AQ4 function in certain regions, it could potentially delay or hinder the proper maturation of the gliovascular in a region-specific manner.

This study reveals that distinct brain regions exhibit varying responses to prenatal stress, with the cerebellum appearing less prone to alterations. In contrast, we identified the hippocampus and prefrontal cortex as particularly vulnerable regions in females exposed to prenatal stress. This observation aligns with prior research in rodent models of prenatal DEX exposure, which identified sex-specific morphological changes in microglia within these regions, potentially contributing to the susceptibility to psychiatric disorders like chronic anxiety and depression.(Caetano et al., 2017; Gaspar et al., 2021). The hippocampus and prefrontal cortex are essential for critical cognitive functions such as learning, memory, planning, decision-making, and working memory (Rubin et al., 2014; Szczepanski & Knight, 2014). Dysfunctions in these processes can lead to behavioral alterations and elevate the risk of developing sex-specific mental disorders later in life (Anderson et al., 2009; Sigurdsson & Duvarci, 2015). Notably, women are reported to have a higher susceptibility to depression and anxiety disorders compared to men (Albert, 2015; McLean et al., 2011). Interestingly, existing data suggests a potential sex difference, with lower female astrocyte density compared to males in response to stress (Orso et al., 2023). This increased vulnerability in females (Goodwill et al., 2019; Wellman et al., 2018) might be associated with heightened astrocytic sensitivity to stress.

Understanding the impact of early life adversities on infancy is crucial, as it can reveal alterations that may allow for early intervention. Here, we break new ground by investigating the response of astrocyte endfeet to prenatal stress, an area that has previously been unexplored. Further research is essential to explore the long-term consequences of these stress-induced alterations and identify potential interventions to mitigate their effects.

## References

Albert, P. R. (2015). Why is depression more prevalent in women? Journal of Psychiatry & Neuroscience : JPN, 40(4), 219–221. 10.1503/jpn.150205

Anderson, S. W., Wisnowski, J. L., Barrash, J., Damasio, H., & Tranel, D. (2009). Consistency of neuropsychological outcome following damage to prefrontal cortex in the first years of life. Journal of Clinical and Experimental Neuropsychology, 31(2), 170–179. 10.1080/13803390802360526

Bock, J., Wainstock, T., Braun, K., & Segal, M. (2015). Stress In Utero: Prenatal Programming of Brain Plasticity and Cognition. Biological Psychiatry, 78(5), 315–326. 10.1016/j.biopsych.2015.02.036

Bolte, S., & Cordelières, F. P. (2006). A guided tour into subcellular colocalization analysis in light microscopy. Journal of Microscopy, 224(Pt 3), 213–232. 10.1111/j.1365-2818.2006.01706.x

Borges, S., Coimbra, B., Soares-Cunha, C., Miguel Pêgo, J., Sousa, N., & João Rodrigues, A. (2013). Dopaminergic modulation of affective and social deficits induced by prenatal glucocorticoid exposure. Neuropsychopharmacology : Official Publication of the American College of Neuropsychopharmacology, 38(10), 2068–2079. 10.1038/npp.2013.108

Caetano, L., Pinheiro, H., Patrício, P., Mateus-Pinheiro, A., Alves, N. D., Coimbra, B., Baptista, F. I., Henriques, S. N., Cunha, C., Santos, A. R., Ferreira, S. G., Sardinha, V. M., Oliveira, J. F., Ambrósio, A. F., Sousa, N., Cunha, R. A., Rodrigues, A. J., Pinto, L., & Gomes, C. A. (2017). Adenosine A2A receptor regulation of microglia morphological remodeling-gender bias in physiology and in a model of chronic anxiety. Molecular Psychiatry, 22(7), 1035–1043. 10.1038/mp.2016.173

Coelho-Santos, V., & Shih, A. Y. (2020). Postnatal development of cerebrovascular structure and the neurogliovascular unit. Wiley Interdisciplinary Reviews. Developmental Biology, 9(2), e363. 10.1002/wdev.363

Crochemore, C., Lu, J., Wu, Y., Liposits, Z., Sousa, N., Holsboer, F., & Almeida, O. F. X. (2005). Direct targeting of hippocampal neurons for apoptosis by glucocorticoids is reversible by mineralocorticoid receptor activation. Molecular Psychiatry, 10(8), 790–798. 10.1038/sj.mp.4001679

Davis, E. P., Sandman, C. A., Buss, C., Wing, D. A., & Head, K. (2013). Fetal glucocorticoid exposure is associated with preadolescent brain development. Biological Psychiatry, 74(9), 647–655. 10.1016/j.biopsych.2013.03.009

Dion-Albert, L., Cadoret, A., Doney, E., Kaufmann, F. N., Dudek, K. A., Daigle, B., Parise, L. F., Cathomas, F., Samba, N., Hudson, N., Lebel, M., Signature Consortium, Campbell, M., Turecki, G., Mechawar, N., & Menard, C. (2022). Vascular and blood-brain barrier-related changes underlie stress responses and resilience in female mice and depression in human tissue. Nature Communications, 13(1), 164. 10.1038/s41467-021-27604-x

Dion-Albert, L., Dudek, K. A., Russo, S. J., Campbell, M., & Menard, C. (2023). Neurovascular adaptations modulating cognition, mood, and stress responses. Trends in Neurosciences, 46(4), 276–292. 10.1016/j.tins.2023.01.005

Drozdowicz, L. B., & Bostwick, J. M. (2014). Psychiatric adverse effects of pediatric corticosteroid use. Mayo Clinic Proceedings, 89(6), 817–834. 10.1016/j.mayocp.2014.01.010

Dudek, K. A., Dion-Albert, L., Lebel, M., LeClair, K., Labrecque, S., Tuck, E., Ferrer Perez, C., Golden, S. A., Tamminga, C., Turecki, G., Mechawar, N., Russo, S. J., & Menard, C. (2020). Molecular adaptations of the blood-brain barrier promote stress resilience vs. depression. Proceedings of the National Academy of Sciences of the United States of America, 117(6), 3326–3336. 10.1073/pnas.1914655117

Fallier-Becker, P., Vollmer, J. P., Bauer, H.-C., Noell, S., Wolburg, H., & Mack, A. F. (2014). Onset of aquaporin-4 expression in the developing mouse brain. International Journal of Developmental Neuroscience : The Official Journal of the International Society for Developmental Neuroscience, 36, 81–89. 10.1016/j.ijdevneu.2014.06.001

Freitas-Andrade, M., Comin, C. H., Van Dyken, P., Ouellette, J., Raman-Nair, J., Blakeley, N., Liu, Q. Y., Leclerc, S., Pan, Y., Liu, Z., Carrier, M., Thakur, K., Savard, A., Rurak, G. M., Tremblay, M.-È., Salmaso, N., da F Costa, L., Coppola, G., & Lacoste, B. (2023). Astroglial Hmgb1 regulates postnatal astrocyte morphogenesis and cerebrovascular maturation. Nature Communications, 14(1), 4965. 10.1038/s41467-023-40682-3

Fukumoto, K., Morita, T., Mayanagi, T., Tanokashira, D., Yoshida, T., Sakai, A., & Sobue, K. (2009). Detrimental effects of glucocorticoids on neuronal migration during brain development. Molecular Psychiatry, 14(12), 1119–1131. 10.1038/mp.2009.60

Gaspar, R., Soares-Cunha, C., Domingues, A. V., Coimbra, B., Baptista, F. I., Pinto, L., Ambrósio, A. F., Rodrigues, A. J., & Gomes, C. A. (2021). Resilience to stress and sex-specific remodeling of microglia and neuronal morphology in a rat model of anxiety and anhedonia. Neurobiology of Stress, 14, 100302. 10.1016/j.ynstr.2021.100302

Gilbert, A., Vidal, X. E., Estevez, R., Cohen-Salmon, M., & Boulay, A.-C. (2019). Postnatal development of the astrocyte perivascular MLC1/GlialCAM complex defines a temporal window for the gliovascular unit maturation. Brain Structure & Function, 224(3), 1267–1278. 10.1007/s00429-019-01832-w

Goodwill, H. L., Manzano-Nieves, G., Gallo, M., Lee, H.-I., Oyerinde, E., Serre, T., & Bath, K. G. (2019). Early life stress leads to sex differences in development of depressive-like outcomes in a mouse model. Neuropsychopharmacology : Official Publication of the American College of Neuropsychopharmacology, 44(4), 711–720. 10.1038/s41386-018-0195-5

Haj-Yasein, N. N., Jensen, V., Østby, I., Omholt, S. W., Voipio, J., Kaila, K., Ottersen, O. P., Hvalby, Ø., & Nagelhus, E. A. (2012). Aquaporin-4 regulates extracellular space volume dynamics during high-frequency synaptic stimulation: a gene deletion study in mouse hippocampus. Glia, 60(6), 867–874. 10.1002/glia.22319

Han, H.-C. (2012). Twisted blood vessels: symptoms, etiology and biomechanical mechanisms. Journal of Vascular Research, 49(3), 185–197. 10.1159/000335123

Igarashi, H., Tsujita, M., Kwee, I. L., & Nakada, T. (2014). Water influx into cerebrospinal fluid is primarily controlled by aquaporin-4, not by aquaporin-1: 17O JJVCPE MRI study in knockout mice. Neuroreport, 25(1), 39–43. 10.1097/WNR.0000000000000042

Khulan, B., & Drake, A. J. (2012). Glucocorticoids as mediators of developmental programming effects. Best Practice & Research. Clinical Endocrinology & Metabolism, 26(5), 689–700. 10.1016/j.beem.2012.03.007

Krontira, A. C., Cruceanu, C., & Binder, E. B. (2020). Glucocorticoids as Mediators of Adverse Outcomes of Prenatal Stress. Trends in Neurosciences, 43(6), 394–405. 10.1016/j.tins.2020.03.008

Li, X., Gao, J., Ding, J., Hu, G., & Xiao, M. (2013). Aquaporin-4 expression contributes to decreases in brain water content during mouse postnatal development. Brain Research Bulletin, 94, 49–55. 10.1016/j.brainresbull.2013.02.004

Lunde, L. K., Camassa, L. M. A., Hoddevik, E. H., Khan, F. H., Ottersen, O. P., Boldt, H. B., & Amiry-Moghaddam, M. (2015). Postnatal development of the molecular complex underlying astrocyte polarization. Brain Structure & Function, 220(4), 2087–2101. 10.1007/s00429-014-0775-z

Majcher-Maślanka, I., Solarz, A., & Chocyk, A. (2019). Maternal separation disturbs postnatal development of the medial prefrontal cortex and affects the number of neurons and glial cells in adolescent rats. Neuroscience, 423, 131–147. 10.1016/j.neuroscience.2019.10.033

Martisova, E., Aisa, B., Guereñu, G., & Ramírez, M. J. (2013). Effects of early maternal separation on biobehavioral and neuropathological markers of Alzheimer’s disease in adult male rats. Current Alzheimer Research, 10(4), 420–432. 10.2174/1567205011310040007

McLean, C. P., Asnaani, A., Litz, B. T., & Hofmann, S. G. (2011). Gender differences in anxiety disorders: prevalence, course of illness, comorbidity and burden of illness. Journal of Psychiatric Research, 45(8), 1027–1035. 10.1016/j.jpsychires.2011.03.006

Millage, A. R., Latuga, M. S., & Aschner, J. L. (2017). Effect of perinatal glucocorticoids on vascular health and disease. Pediatric Research, 81(1–1), 4–10. 10.1038/pr.2016.188

Najjar, S., Pahlajani, S., De Sanctis, V., Stern, J. N. H., Najjar, A., & Chong, D. (2017). Neurovascular Unit Dysfunction and Blood-Brain Barrier Hyperpermeability Contribute to Schizophrenia Neurobiology: A Theoretical Integration of Clinical and Experimental Evidence. Frontiers in Psychiatry, 8, 83. 10.3389/fpsyt.2017.00083

Nakada, T., Kwee, I. L., Igarashi, H., & Suzuki, Y. (2017). Aquaporin-4 Functionality and Virchow-Robin Space Water Dynamics: Physiological Model for Neurovascular Coupling and Glymphatic Flow. International Journal of Molecular Sciences, 18(8). 10.3390/ijms18081798

Neigh, G. N., Owens, M. J., Taylor, W. R., & Nemeroff, C. B. (2010). Changes in the vascular area fraction of the hippocampus and amygdala are induced by prenatal dexamethasone and/or adult stress. Journal of Cerebral Blood Flow and Metabolism : Official Journal of the International Society of Cerebral Blood Flow and Metabolism, 30(6), 1100–1104. 10.1038/jcbfm.2010.46

Neuhaus, W., Schlundt, M., Fehrholz, M., Ehrke, A., Kunzmann, S., Liebner, S., Speer, C. P., & Förster, C. Y. (2015). Multiple Antenatal Dexamethasone Treatment Alters Brain Vessel Differentiation in Newborn Mouse Pups. PloS One, 10(8), e0136221. 10.1371/journal.pone.0136221

Oliveira, M., Bessa, J. M., Mesquita, A., Tavares, H., Carvalho, A., Silva, R., Pêgo, J. M., Cerqueira, J. J., Palha, J. A., Almeida, O. F. X., & Sousa, N. (2006). Induction of a hyperanxious state by antenatal dexamethasone: a case for less detrimental natural corticosteroids. Biological Psychiatry, 59(9), 844– 852. 10.1016/j.biopsych.2005.08.020

Oliveira, M., Rodrigues, A.-J., Leão, P., Cardona, D., Pêgo, J. M., & Sousa, N. (2012). The bed nucleus of stria terminalis and the amygdala as targets of antenatal glucocorticoids: implications for fear and anxiety responses. Psychopharmacology, 220(3), 443–453. 10.1007/s00213-011-2494-y

Orso, R., Creutzberg, K. C., Lumertz, F. S., Kestering-Ferreira, E., Stocchero, B. A., Perrone, M. K., Begni, V., Grassi-Oliveira, R., Riva, M. A., & Viola, T. W. (2023). A systematic review and multilevel meta-analysis of the prenatal and early life stress effects on rodent microglia, astrocyte, and oligodendrocyte density and morphology. Neuroscience and Biobehavioral Reviews, 150, 105202. 10.1016/j.neubiorev.2023.105202

Ouellette, J., Toussay, X., Comin, C. H., Costa, L. da F., Ho, M., Lacalle-Aurioles, M., Freitas-Andrade, M., Liu, Q. Y., Leclerc, S., Pan, Y., Liu, Z., Thibodeau, J.-F., Yin, M., Carrier, M., Morse, C. J., Dyken, P. Van, Bergin, C. J., Baillet, S., Kennedy, C. R., … Lacoste, B. (2020). Vascular contributions to 16p11.2 deletion autism syndrome modeled in mice. Nature Neuroscience, 23(9), 1090–1101. 10.1038/s41593-020-0663-1

Pinheiro, H., Gaspar, R., Baptista, F. I., Fontes-Ribeiro, C. A., Ambrósio, A. F., & Gomes, C. A. (2018). Adenosine A2A Receptor Blockade Modulates Glucocorticoid-Induced Morphological Alterations in Axons, But Not in Dendrites, of Hippocampal Neurons. Frontiers in Pharmacology, 9, 219. 10.3389/fphar.2018.00219

Polytarchou, C., & Papadimitriou, E. (2005). Antioxidants inhibit human endothelial cell functions through down-regulation of endothelial nitric oxide synthase activity. European Journal of Pharmacology, 510(1–2), 31–38. 10.1016/j.ejphar.2005.01.004

Rim, C., Park, H.-S., You, M.-J., Yang, B., Kim, H.-J., Sung, S., & Kwon, M.-S. (2022). Microglia involvement in sex-dependent behaviors and schizophrenia occurrence in offspring with maternal dexamethasone exposure. Schizophrenia (Heidelberg, Germany), 8(1), 71. 10.1038/s41537-022-00280-6

Rodrigues, A. J., Leão, P., Pêgo, J. M., Cardona, D., Carvalho, M. M., Oliveira, M., Costa, B. M., Carvalho, A. F., Morgado, P., Araújo, D., Palha, J. A., Almeida, O. F. X., & Sousa, N. (2012). Mechanisms of initiation and reversal of drug-seeking behavior induced by prenatal exposure to glucocorticoids. Molecular Psychiatry, 17(12), 1295–1305. 10.1038/mp.2011.126

Rubin, R. D., Watson, P. D., Duff, M. C., & Cohen, N. J. (2014). The role of the hippocampus in flexible cognition and social behavior. Frontiers in Human Neuroscience, 8, 742. 10.3389/fnhum.2014.00742

Sadowska, G. B., Malaeb, S. N., & Stonestreet, B. S. (2010). Maternal glucocorticoid exposure alters tight junction protein expression in the brain of fetal sheep. American Journal of Physiology. Heart and Circulatory Physiology, 298(1), H179–88. 10.1152/ajpheart.00828.2009

Salman, M. M., Kitchen, P., Halsey, A., Wang, M. X., Törnroth-Horsefield, S., Conner, A. C., Badaut, J., Iliff, J. J., & Bill, R. M. (2022). Emerging roles for dynamic aquaporin-4 subcellular relocalization in CNS water homeostasis. Brain : A Journal of Neurology, 145(1), 64–75. 10.1093/brain/awab311

Shende, V. H., McArthur, S., Gillies, G. E., & Opacka-Juffry, J. (2015). Astroglial Plasticity Is Implicated in Hippocampal Remodelling in Adult Rats Exposed to Antenatal Dexamethasone. Neural Plasticity, 2015, 694347. 10.1155/2015/694347

Sigurdsson, T., & Duvarci, S. (2015). Hippocampal-Prefrontal Interactions in Cognition, Behavior and Psychiatric Disease. Frontiers in Systems Neuroscience, 9, 190. 10.3389/fnsys.2015.00190

Szczepanski, S. M., & Knight, R. T. (2014). Insights into human behavior from lesions to the prefrontal cortex. Neuron, 83(5), 1002–1018. 10.1016/j.neuron.2014.08.011

Tanokashira, D., Morita, T., Hayashi, K., Mayanagi, T., Fukumoto, K., Kubota, Y., Yamashita, T., & Sobue, K. (2012). Glucocorticoid suppresses dendritic spine development mediated by down-regulation of caldesmon expression. The Journal of Neuroscience : The Official Journal of the Society for Neuroscience, 32(42), 14583–14591. 10.1523/JNEUROSCI.2380-12.2012

Tsiarli, M. A., Rudine, A., Kendall, N., Pratt, M. O., Krall, R., Thiels, E., DeFranco, D. B., & Monaghan, A. P. (2017). Antenatal dexamethasone exposure differentially affects distinct cortical neural progenitor cells and triggers long-term changes in murine cerebral architecture and behavior. Translational Psychiatry, 7(6), e1153. 10.1038/tp.2017.65

Vivi, E., & Di Benedetto, B. (2024). Brain stars take the lead during critical periods of early postnatal brain development: relevance of astrocytes in health and mental disorders. Molecular Psychiatry. 10.1038/s41380-024-02534-4

Wallerath, T., Gödecke, A., Molojavyi, A., Li, H., Schrader, J., & Förstermann, U. (2004). Dexamethasone lacks effect on blood pressure in mice with a disrupted endothelial NO synthase gene. Nitric Oxide : Biology and Chemistry, 10(1), 36–41. 10.1016/j.niox.2004.01.008

Wellman, C. L., Bangasser, D. A., Bollinger, J. L., Coutellier, L., Logrip, M. L., Moench, K. M., & Urban, K. R. (2018). Sex Differences in Risk and Resilience: Stress Effects on the Neural Substrates of Emotion and Motivation. The Journal of Neuroscience : The Official Journal of the Society for Neuroscience, 38(44), 9423–9432. 10.1523/JNEUROSCI.1673-18.2018

Wen, H., Nagelhus, E. A., Amiry-Moghaddam, M., Agre, P., Ottersen, O. P., & Nielsen, S. (1999). Ontogeny of water transport in rat brain: postnatal expression of the aquaporin-4 water channel. The European Journal of Neuroscience, 11(3), 935–945. 10.1046/j.1460-9568.1999.00502.x

Zeng, H., Zhang, X., Wang, W., Shen, Z., Dai, Z., Yu, Z., Xu, S., Yan, G., Huang, Q., Wu, R., Chen, X., & Xu, H. (2020). Maternal separation with early weaning impairs neuron-glia integrity: non-invasive evaluation and substructure demonstration. Scientific Reports, 10(1), 19440. 10.1038/s41598-020-76640-y

